# Cryo-EM structure of cardiac amyloid fibrils from an immunoglobulin light chain (AL) amyloidosis patient

**DOI:** 10.1101/444901

**Authors:** Paolo Swuec, Francesca Lavatelli, Masayoshi Tasaki, Cristina Paissoni, Paola Rognoni, Martina Maritan, Francesca Brambilla, Paolo Milani, Pierluigi Mauri, Carlo Camilloni, Giovanni Palladini, Giampaolo Merlini, Stefano Ricagno, Martino Bolognesi

## Abstract

Systemic light chain (AL) amyloidosis is a life-threatening disease caused by aggregation and deposition of monoclonal immunoglobulin light chains (LC) in target organs. Severity of heart involvement is the most important factor determining prognosis. Here, we report the 4.0 Å resolution cryo-electron microscopy (cryo-EM) map and structural model of amyloid fibrils extracted from the heart of an AL patient affected by severe amyloid cardiomyopathy. The fibrils are composed of one asymmetric protofilament, showing typical 4.9 Å stacking and parallel cross-β architecture. Two distinct polypeptide stretches belonging to the LC variable domain (V_l_) could be modelled in the density (total of 77 residues), stressing the role of the V_l_ domain in fibril assembly and LC aggregation. Despite high levels of V_l_ sequence variability, residues stabilising the observed fibril core are conserved through several V_l_ domains, highlighting structural motifs that may be common to misfolded LCs. Our data shed first light on the architecture of life-threatening LC amyloid deposits, and provide a rationale for correlating LC amino acid sequences and fibril structures.

---

Light chain (AL) amyloidosis, with an incidence of about 10 new cases per million-persons/year, is currently the most common systemic form of amyloidosis in Western countries^1^. The disease is associated with the presence of a plasma cell clone, and is caused by extracellular deposition of misfolding-prone monoclonal immunoglobulin light chains (LCs), transported to target organs through blood. Deposition of amyloid fibrils is associated with dysfunction of affected organs. The amino acid sequence of each patient’s monoclonal LC is virtually unique, as a consequence of immunoglobulin germline genes rearrangement and somatic hypermutation. Fibril deposition in AL is widespread, and can target different organs; heart involvement dramatically worsens patients’ prognosis^2-4^. Much research is currently being devoted to defining the molecular bases of amyloid cardiomyopathy^5-7^, to hinder fibrillogenesis^8^ and cell damage^5,9,10^.

LC subunits (*ca.* 215 residues) consist of two β-sandwich domains, each hosting a disulphide bridge: the highly variable N-terminal domain (V_l_; *ca.* 110 residues), a short joining region (J_l_), and the C-terminal constant domain (C_l_)^6,11^. Both full-length LCs and isolated V_l_ domains are typical components of the deposited fibrils^12,13^; nonetheless, the mechanisms promoting aggregation *in vivo* remain unclear. Progress in understanding LC aggregation is hampered by lack of structural insight on AL fibrils, only low-resolution characterisation of LC fibrils being available to date^14,15^.

Cryo-EM proved as a first-choice method for structural analyses of amyloid aggregates prepared *in vitro* or extracted *ex vivo*^16-20^. Notably, in the few reported cases, the fibril component protein was shown to adopt complex folds, compatible but not fully predictable from fibril models based on short peptides^21^. Here we report the cryo-EM structure, at overall resolution of 4.0 Å, of *ex vivo* heart LC fibrils from a patient affected by severe cardiac amyloidosis.

In order to define the structure of natural amyloid fibrils, we extracted and characterized *ex vivo* amyloid aggregates from the affected heart tissue. Specifically, fibrils were isolated from left ventricle specimens acquired during autopsy from a male patient affected by AL λ amyloidosis, with severe amyloid cardiomyopathy (**Fig.** 1A and Extended Data **Table** 1). The monoclonal amyloidogenic LC, labelled as AL55, was sequenced from its coding mRNA from bone marrow plasma cells; AL55 is of λ isotype and belongs to the *IGLV6-57* germline gene.

**Figure 1.**
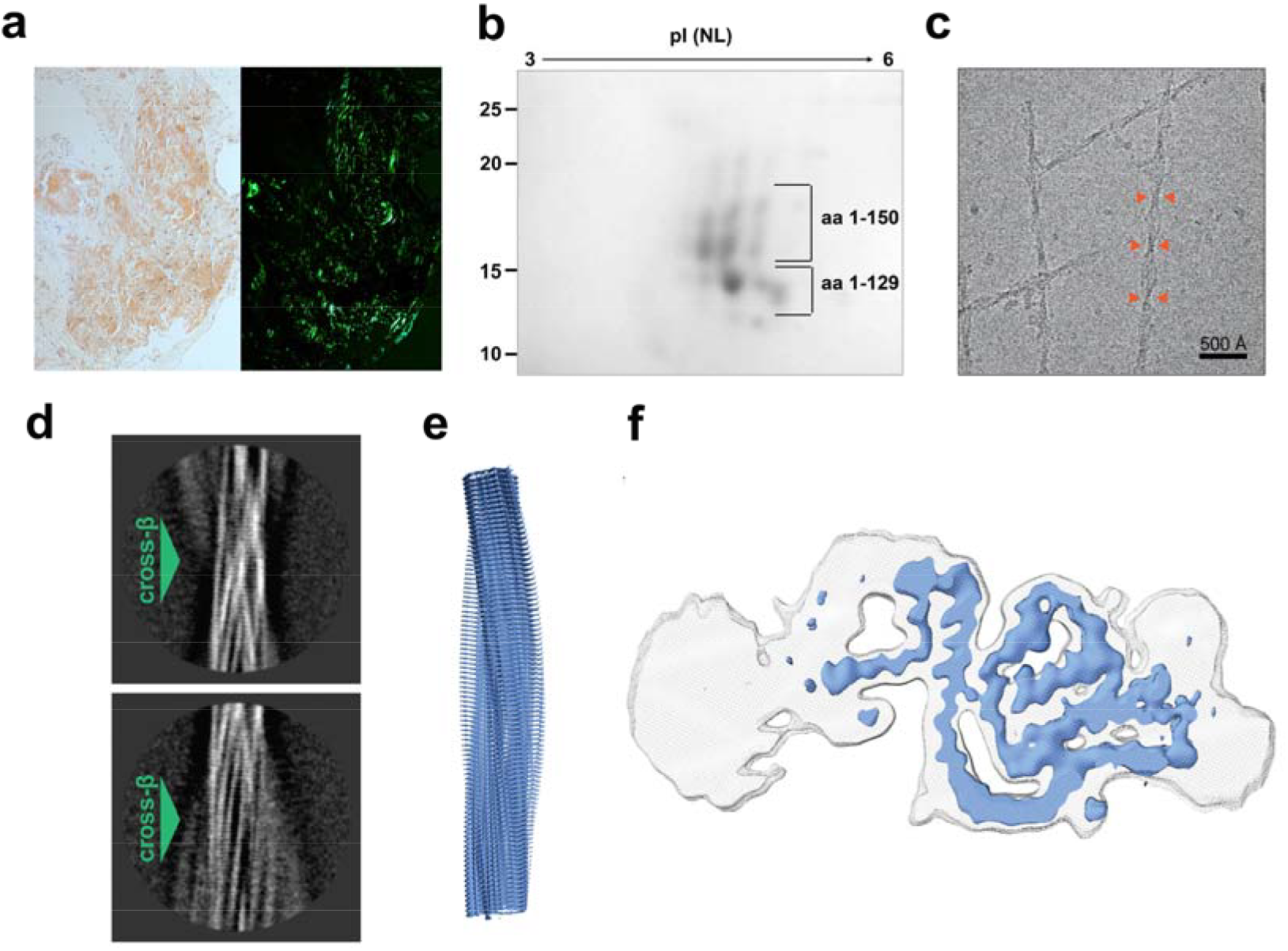
Structure of AL55 fibrils from heart tissue. **a**, Myocardial tissue from patient AL55, stained with Congo red. Red-orange stain and apple-green birefringence indicate amyloid deposits under visible (left) and under polarized light (right), respectively. **b**, 2D-PAGE analysis of purified AL55 LC fibrils (inset from Extended Data **Fig**. 1b, left panel). Low and high MW fragments comprise residues from the N-*terminus* to residue 129, and to 150, respectively. **c**, Representative cryo-EM micrograph of AL55 LC fibrils; orange arrows highlight fibril cross-overs. **d**, Reference-free 2D class averages of AL55 fibril showing distinct cross-β staggering (green arrows). **e**, Overview of the AL55 fibril cryo-EM map covering the structured fibril core region (blue). **f**, Cross-sectional EM densities of sharpened, 4.0 Å (blue) and unsharpened, 4.5 Å low-pass filtered (grey) reconstructions.

The molecular composition of the isolated fibrils was analysed by a proteomic approach, based on two-dimensional polyacrylamide gel electrophoresis (2D-PAGE) coupled to nano-liquid chromatography tandem mass spectrometry (nLC-MS/MS). In agreement with previous observations ^12,13^, fibrils are composed of a heterogeneous population of LC proteoforms and N-terminal fragments (**Fig.** 1b and Extended Data **Fig**. 1b). The LC fragments with lower molecular weight are predominant, and mainly contain the V_l_ domain. However, longer fragments, extending through the constant domain, and species consistent with the full length LC are also present. Low molecular weight fragments are resistant to limited proteolysis (Extended Data Fig. 1b), whereas degradation of the longer proteoforms is observed, suggesting protection or burial of the V_l_ domain within the fibril.

Contrary to previous reports^14^, our negative staining electron microscopy and cryo-EM analyses of freshly extracted amyloid material revealed no polymorphs (**Fig**. 1c and Extended Data **Fig**. 2). AL55 fibrils display a helical pitch of 1070 ± 30 Å, and 80 - 175 Å width range (Extended Data **Fig**. 2). Absence of two symmetric protofilaments is evident from inspection of the fibrils in raw micrographs and in images obtained by reference-free 2D classification of segments comprising an entire helical pitch (**Fig**. 1c and Extended Data **Fig**. 2). 2D class averages of vitrified AL55 fibrils, where β-strands are clearly resolved, suggest the presence of a highly ordered core surrounded by low order regions (**Fig**. 1d and Extended Data **Fig**. 2).

Cryo-EM 3D reconstruction of AL55 fibrils resulted in a map at overall resolution of 4.0 Å, in which the cross-β structure was clearly resolved (**Fig**. 1e-f and Extended Data **Fig**. 3). Consistent with our 2D analyses, the AL55 map showed a core whose consecutive β-strand rungs are related by helical symmetry, with a rise of 4.9 Å, a twist of approximately – 1.6°, and two low-order outer regions.

For each LC subunit deposited along the fibril axis, the structured core is divided into two segments: the central part of the density displays a ‘snail-shell’ trace that is surrounded by a second, ‘C-shaped’, extended polypeptide stretch (**Fig.** 1f). The two regions are spatially contiguous but not directly connected by interpretable density, indicating that two distinct LC segments build the fibril core. Several bulky side-chains visible in the map, together with the C22-C91 disulphide bridge, supported chain tracing and modelling for 77 residues of AL55 V_l_. As a result, the first N-terminal 37 residues map into the internal snail-shell region, while the outer C-shaped stretch hosts residues 66-105 (**Fig.** 2a-b).

**Figure 2.**
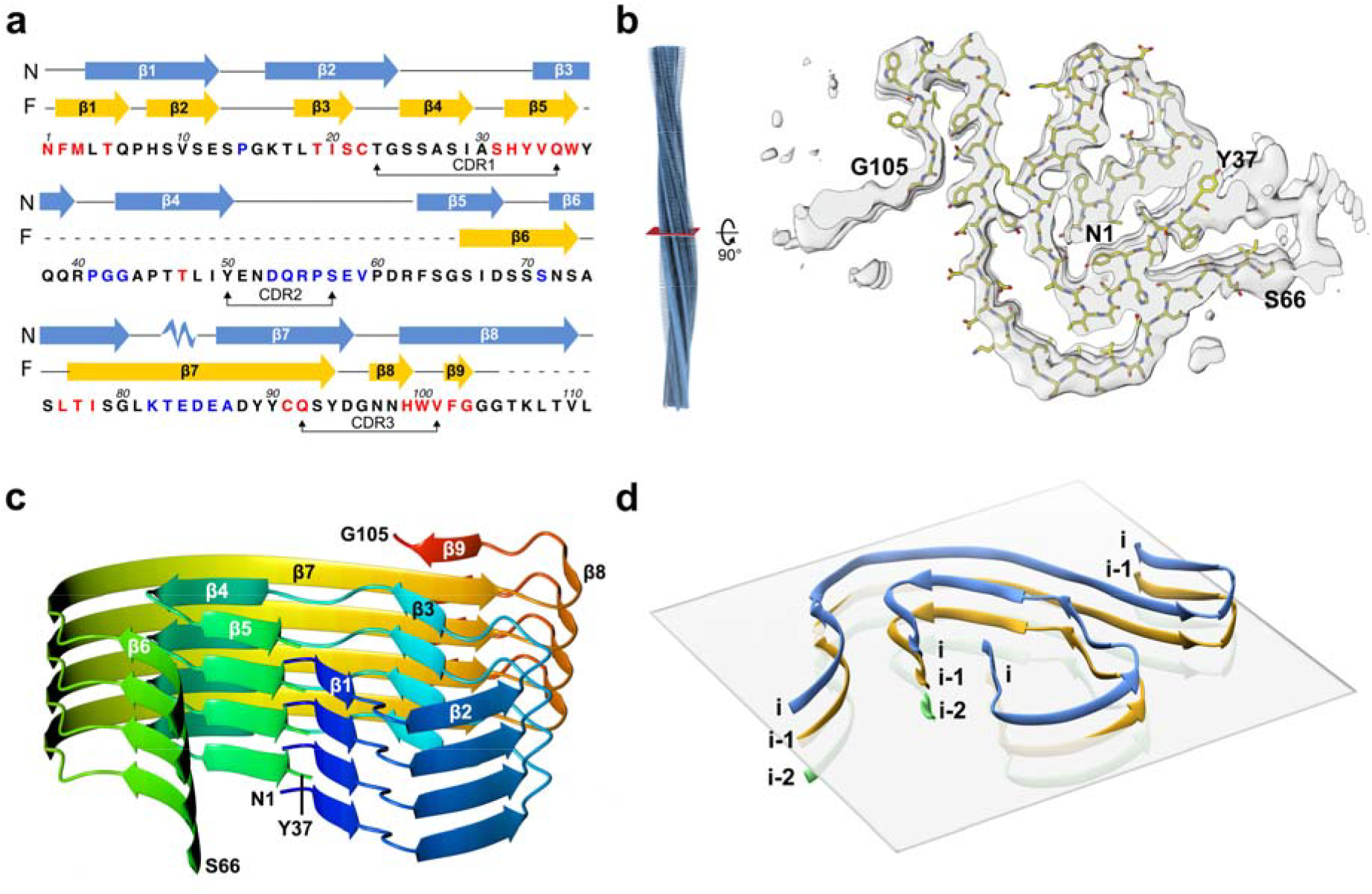
V_l_ domain structured fragment in the fibril core. **a**, Sequence of AL55 V_l_ domain: scheme of the secondary structure elements of the natively folded LC (N, blue), and of the fibril assembly (F, yellow), respectively; dashed lines correspond to residues non-modelled in the EM density. Residues are coloured according to their intrinsic aggregation propensities, as defined by CamSol^22^ (high, in red; low, in black; very low in blue). **b**, Overview of AL55 fibril helical reconstruction (left) and cross-sectional densities (right). Atomic model of AL55 (residues 1-37 and 66-105) superimposed on a cross-section of the EM density map. **c**, Ribbon representation of the fibril structured core, rainbow coloured. Four stacks (subunits) in the typical cross-β arrangement are shown. **d**, A horizontal plane (perpendicular to the fibril elongation axis) is added in this ribbon representation of the fibril core to highlight the raise of each subunit at the end of β2 (P15) and of β5 (W36), respectively.

Individual LC subunits assemble with a parallel β-sheet topology along the fibril elongation axis, *i.e.* along the inter-subunit H-bonding direction. Each subunit presents nine β-strands; β1-β5 belong to the snail-shell region, and β6-β9 pack around it in the C-shaped stretch (**Fig.** 2c and 3a). In particular, β1, β3, β5 and β6 face each other and tightly pack their side chains together, while β4, β7 and β9 form a second contact region of lower side chain packing density.

**Figure 3.**
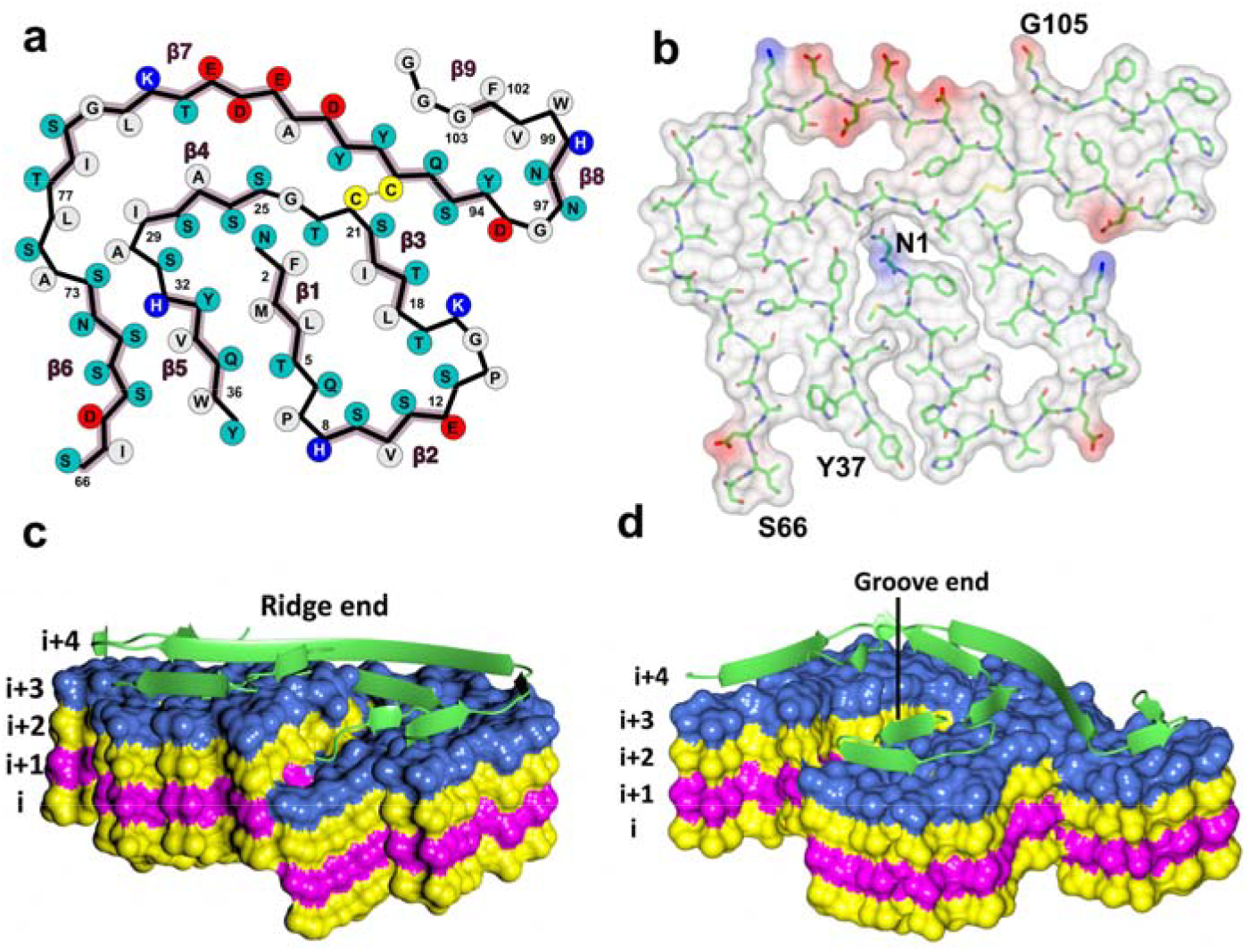
Interactions and interfaces in the fibril core. **a**, A 2D schematic view of the core polypeptide stretches. Residues are coloured as follows: grey hydrophobic, cyan polar, red and blue negatively and positively charged, respectively. **b**, Stick representation of the fibril core together with a surface representation coloured according to electrostatic charges. **c-d**, A structural view of the fibril ends: the inner core (β1- β2) of the fibril is shown protruding from the surface (ridge; **c**); inner core concave region (groove) is shown in **d**.

As previously reported for other amyloid fibril structures^16-18,20,23^, the β1-β5 modelled fragment does not display a planar arrangement in the fibril, the snail shell region adopting a β-helix-like structure. In particular, the polypeptide chain of subunit *i* rises twice, at residues P14 (*i*+1) and at W36 (*i*+2), such that the side chains from β1 of subunit *i* pack against the C-terminal residues of subunit *i*-2 (**Fig.** 2d). On the contrary, the C-shaped region (β6-β9) lays essentially in a plane; given that it is covalently bound to the snail shell region through the C22-C91 disulphide bond, this segment in molecule *i* is located at *i*+1 level (**Fig.** 2d). Such overall assembly produces non-planar fibril ends. Analogously to what has been discussed for amyloid-β (1-42) fibrils^18^, the two ends of a fibril present a groove and a ridge (**Fig.** 3c-d), both of which expose highly hydrophobic patches (β1- β3 interface). Conceivably, the edge strand β1 of a natively folded LC could be recruited by the hydrophobic groove/ridge of a growing fibril, promoting unfolding and association of a new subunit.

Overall the AL55 segments that build the rigid fibril core are characterised by a few charged residues (9 out of 77), four of which are involved in two salt bridges (K16-D95, K82-E84) alleviating Coulombic repulsion (**Fig.** 3b). The 82-88 segment, which comprises most of the charged residues, is detached from the inner fibril core (**Fig.** 3b) and may prove accessible to water molecules. The V_l_ stretches predicted to be most aggregation prone^22^ are all comprised in the fibril core (**Fig.** 2a) that hosts two hydrophobic clusters. The first cluster builds the β1 - β3 interface (F2, L4, L18, I20), while the second is located between the β4-β5 turn and β6-β7 (I29, L76, I78, L81) (**Fig.** 3a), and includes the disulphide bond linking β4 to β7. The two Pro residues in the fibril core (P7 and P14) help forming the β1-β2 and β2-β3 turns, respectively. In the native V_l_ immunoglobulin fold P7 is located at the centre of the edge β1-strand that is bulged; notably, such bulged edge strands are held to protect against amyloid aggregation^24^. Nevertheless, in the AL55 fibril core P7 conveniently accommodates in the β-β2 turn.

Besides the core region, AL55 fibrils present two areas of lower order: the first falls between residues 37 and 66 of the structured segments (**Fig.** 1f and 2b). As limited proteolysis suggests that the whole V_l_ domain is hosted in the fibril core, we postulate that the 38-65 polypeptide fills the low order region adjacent to residues 37 and 66. The 38-65 sequence hosts residues with low β propensity (four Pro and three Gly) and seven charged residues, which might prevent the formation of regular β-structures. The second poorly structured zone surrounds the last modelled residue (G105) (**Fig.** 1f). Mass spectrometry analysis shows that the fibrils contain also portions of the C_l_ domain (that would follow beyond G105 in the intact LC). Such C_l_ portions, which host residues with low aggregation propensity, would fill the poorly defined density region next to the C-end of the modelled polypeptide.

The LC fibril model, here presented, shows that two AL55 hypervariable complementarity determining regions (CDR) build the structured fibril core. CDR1 (T23-Q35) spans β3 through β4; CDR3 (Q92-V101) falls at the C-end of the modelled fibril core. Such observations highlight sequence variability of V_l_ domains as a key factor not only for protein instability and aggregation propensity, but also for the molecular interactions stabilising fibril assembly. Thus, questions arise on how common such fibril architecture might be for AL deposits due to LCs other than AL55. The following structural considerations can be drawn.

Firstly, AL55 solubilised fibrils provide a typical pattern in 2D-PAGE: the fragments of lower molecular weight and higher abundance consist just of the V_l_ domain; longer fragments always contain the LC N-terminal region^13^, strongly suggesting that the V_l_ domain is required to build the fibril core. To assess whether other LCs may be compatible with the fibril architecture here reported, AL55 sequence was aligned with eight diverse cardiotoxic λ LCs (Extended Data **Fig.** 4) previously reported^6^. Despite sequence variability, the residues that appear to play structuring roles in AL55 fibrils are conserved or conservatively mutated. A N-terminal stretch of hydrophobic residues, building the fibril inner β1, is present in all sequences (**Fig**. 3 and Extended Data **Fig**. 4). In several LCs an extra Pro residue (P8) can be found, which would be located in the β1-β2 turn. G15, which in the AL55 fibril core adopts a conformation unfavourable for other amino acids, is conserved in all sequences. The conserved disulphide bond is a strong structural restrain. The two hydrophobic clusters stabilizing the AL55 fibrils can be achieved in all eight λ LCs. Finally, several aromatic residues that stabilise the fibril through inter-subunit stacking interactions, and the (82-88) segment comprising charged residues, are present in all sequences (**Fig**. 3 and Extended Data **Fig**. 4). As observed for two Tau protein isoforms that display different fibril subunit folds^16,17^, sequence variability of the V_l_ domains might also result in some fibril structural rearrangements. Nevertheless, our cryo-EM results on *ex-vivo* cardiac amyloids suggest that structural motifs in the AL55 fibril architecture are compatible with the assembly of amyloid deposit from different LCs, and possibly in other organs, opening the way to a deeper molecular characterisation of AL amyloidosis.

## Data availability

Final AL55 cryo-EM map has been deposited to the Electron Microscopy Data Bank (code EMD-0274). Refined atomic model has been deposited to the Protein Data Bank (code 6HUD). The datasets generated and/or analysed during the current study are available from the corresponding authors upon reasonable request.

## Acknowledgements

Contributions from the University of Milano and Fondazione Romeo and Enrica Invernizzi, to founding the cryo-EM facility are gratefully acknowledged. This work was supported by Fondazione Cariplo (grants n. 2015-0591 and 2016-0489); Associazione Italiana per la Ricerca sul Cancro special program “5 per mille” (number 9965); the Italian Ministry of Health (grants RF-2013-02355259 and RF-2016-02361756), Italian Medicines Agency (grant AIFA-2016- 02364602), and E-Rare JTC 2016 grant ReDox. We are grateful to Laura Verga and Gianluca Capello for immuno-electron microscopy imaging.

## Author contributions

P.S., F.L., MT, C.P., P.M., P.R., M.M., F.B., performed the experiments; P.S., F.L., S.R. designed the experiments; P.S., F.L., C.C., P.M. G.P., G.M., M.B., S.R. wrote the paper with contributions from all other Authors.

## Competing interests

The authors declare no competing interests.

